# Enhanced surface nano-analytics of transient biomolecular processes

**DOI:** 10.1101/2022.07.25.501379

**Authors:** Alyssa Miller, Sean Chia, Zenon Toprakcioglu, Tuuli Hakala, Roman Schmid, Yaduo Feng, Tadas Kartanas, Ayaka Kamada, Michele Vendruscolo, Francesco Simone Ruggeri, Tuomas P. J. Knowles

## Abstract

The study of the physical and chemical properties of biomolecules enables the characterisation of fundamental molecular processes and mechanisms in health and disease. Bulk and single-molecule analytical methods provide rich information on biomolecules, but often require high concentrations and sample preparation away from physiologically relevant conditions. Here, we present the development and application of a lab-on-a-chip approach which combines rapid sample preparation, mixing and deposition to integrate with a range of nano-analytical methods in chemistry and biology, providing enhanced sensitivity and single molecule resolution. We demonstrate that this method empowers multidimensional study of heterogenous biomolecular systems in physiological buffers and concentrations over multiple length scales by nanoscopy and vibrational spectroscopy. We illustrate the capabilities of this platform by capturing and analysing the structural conformations of transient oligomeric species formed at the early stages of the self-assembly of α-synuclein, which are associated with the onset of Parkinson’s disease.

**TEASER:** Maintaining the heterogeneity and structural integrity of monomers and oligomers enables their quantitative study.

## Introduction

Despite significant technological advances in the past few decades, the study of the physical and chemical properties of biomolecules in physiologically relevant solution conditions still represents a significant challenge in the characterisation of biomolecular processes that form the basis of health and disease, as well as for material science applications. Classical bulk analytical approaches such as circular dichroism (CD), mass spectrometry, and vibrational and nuclear magnetic resonance (NMR) spectroscopies, enable chemical identification and structural characterisation. However, they typically require specific sample pre-treatment, or high concentrations and specific buffer conditions which exceed the typical ranges found in functional biological systems.

A solution to these challenges in utilising measurements under bulk conditions is offered by surface-based techniques, which are powerful tools to study the structure and conformation of biomolecules, allowing chemical identification and characterization, such as Raman and infrared (IR) vibrational spectroscopies in attenuated total reflection (ATR-IR), and imaging techniques, such as atomic force microscopy (AFM) and electron microscopy (EM). These well-established surface-based techniques allow for the determination of structural, chemometric and single molecule characterisation of biomolecular processes. However, the deposition of a sample on a surface represents an intrinsic limitation of these surface-methods, because of effects of differential absorption and mass-transport phenomena on the surface^1,2^. For a liquid solution on a surface, the differential interaction and absorption of biomolecules with the surface limits the ability to perform analysisheterogeneous samples, as this selective adsorption results in only a partial representation of the sample composition in bulk solution. This is particularly relevant for single-molecule microscopy and surface-based spectroscopies, since species with potential biological relevance may not be observed, therefore confounding the results^3^. Thus, the solvent is often removed by drying and rinsing of the sample.

Dry conditions also enable analytical measurements which are difficult to conduct in an aqueous environment, such as in the case of nanoscopy and vibrational spectroscopy. However, the removal of the solvent can again introduce artifacts associated with selective adsorption on the surface^1,4^. Furthermore, long incubation times while drying on surfaces can lead to conformational changes, artificial self-assembly, aggregation, and interactions with the surface itself, meaning the conformation in solution is no longer accurately represented^5–11^. This effect can be observed in bulk spectroscopy measurements, which are frequently performed in air to achieve higher sensitivity. It is generally accepted that the drying process may alter the secondary structure of molecules of interest, especially biomolecules that are extremely sensitive to hydration state^2^. These combined limitations prevent the robust assessment of biomolecular structures using bulk and single molecule surface-based analytical methods. Thus, there is an unmet need to overcome the limitations associated with the manual sample preparation for surface-based techniques and allow accurate and reproducible analytical characterisation of heterogenous biomolecular processes.

Here, we report the development of a lab-on-a-chip microfluidic system to overcome the challenges associated with bulk and single-molecule surface analytical chemistry methods on surfaces. This method integrates advances in microfluidic spray deposition with novel lab-on-a-chip capabilities allowing real-time passive mixing, solution changes and dilution of the sample prior to deposition for quantitative nanoscopic, single-molecule and chemometrics analysis^1,12^. We apply this method to study heterogenous biomolecular systems, over multiple biological scales, preserving their conformation in physiological-like buffers and concentrations. We demonstrate that this approach enables the study of samples by several microscopies, including EM, IR spectroscopy and AFM. Novel lab-on-a-chip capabilities allow for on-chip mixing of inorganic, organic and biological samples, for single-step staining and deposition of samples for electron microscopy, thus allowing analysis of morphology while accurately representing sample heterogeneity. Further, we exploit the fast evaporation of droplets generated by the microfluidic spray device to achieve a 100-fold improvement in the sensitivity of commercial vibrational spectroscopy instruments and reach single nanogram protein detection in salt-containing solutions. This procedure enables the preservation and study of the molecular conformations of intrinsically disordered and globular proteins at near-physiological conditions with high sensitivity. Finally, we demonstrate that these capabilities offer highly quantitative information on the conformations of an amyloid-forming protein involved in the onset of Parkinson’s disease and show that we can study the oligomeric species present at the early molecular events of self-assembly, which are generally associated with cellular toxicity.

## Results

### General lab-on-a-chip microfluidic spray deposition

We developed a method with lab-on-a-chip capabilities for sample deposition on surfaces to improve accuracy and reproducibility of a wide range of microscopic and spectroscopic techniques (**Fig. 1**). We innovated the design of our devices to allow passive two solution mixing on-chip, before being intersected with the gas phase to generate droplets^13^ (**Fig. S1**). This allowed for on-chip mixing of solutions and immediate deposition onto the surface with no need for the operator to make changes to the initial experimental set-up (**Fig. S2**).

**Figure 1.**
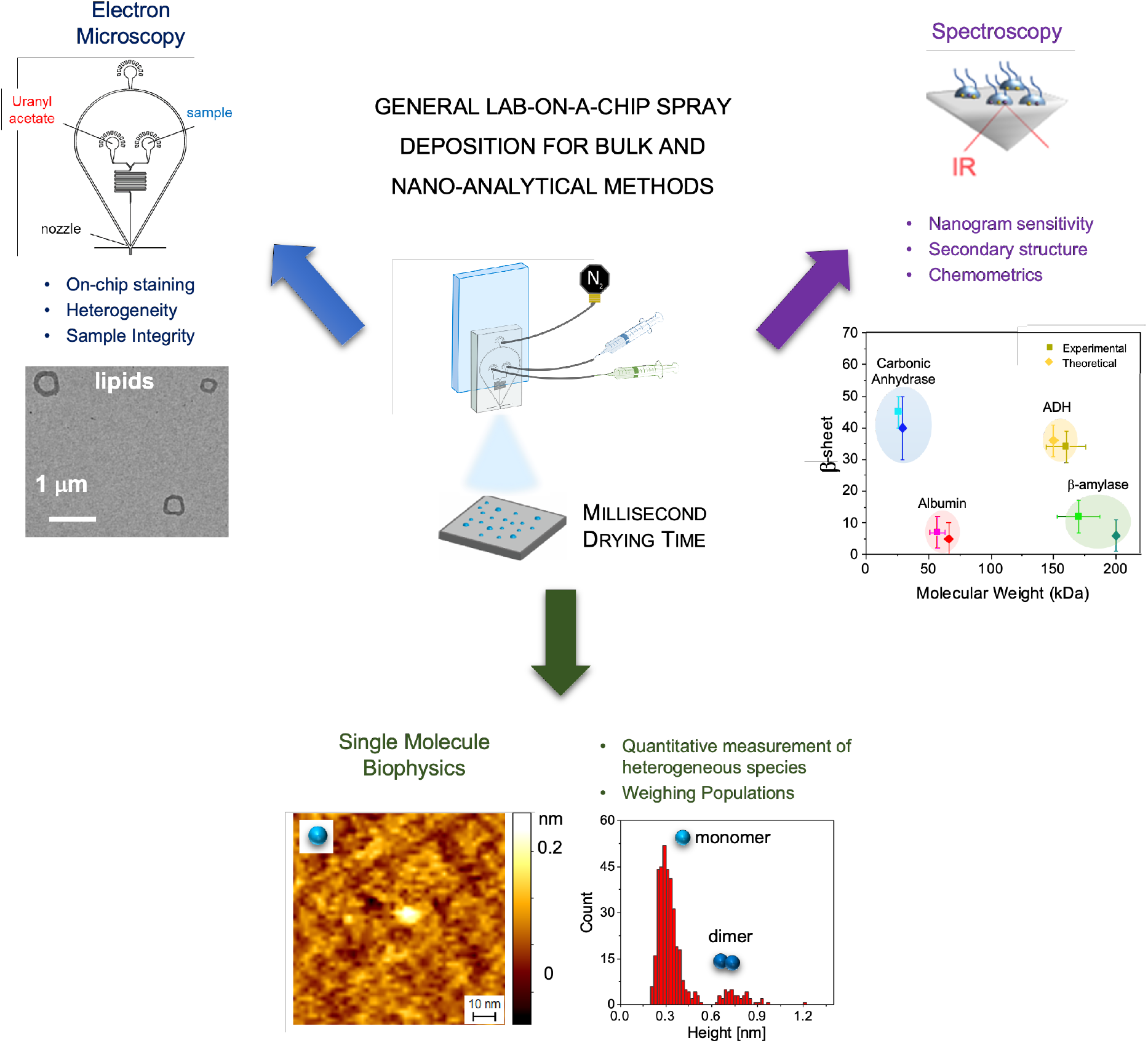
Summary of the applications of the lab-on-a-chip spray deposition method presented in this article. **(a)** In electron microscopy (EM), lab-on-a-chip capabilities are exploited to have on-chip mixing of sample with uranyl acetate stain, allowing for a single-step deposition of a range of materials, while being sufficiently gentle to maintain sample integrity. **(b)** In various spectroscopies, microfluidic spray deposition is applied to the secondary structure and chemometric analysis of proteins in salt-containing solution with unprecedented sensitivity. **(c)** In scanning probe microscopy (SPM), α-synuclein was deposited via microfluidic spray such that the heterogeneity and relative weights of conformational species are represented as they are in solution, thereby enabling their quantitative assessment.

Then, we standardised the spray deposition to ensure high levels of reproducibility and accuracy in the amount and area sprayed between different experiments and operators (**Fig. S3, Supplementary Note 2**). We studied the effect of distance between the device and the surface by varying the distance between 3.5-5.5 cm, as this affects the in-flight drying, and therefore the droplet size which lands on the surface (**Supplementary Note 3**)^12^. We investigated the effect of changing flow rate (50-500 μl/hr) and gas pressure (0.5-4.5 Mbar) to understand their effects on droplet sizes and distributions. This allowed us to determine an optimal, standardised set-up, allowing fine-tuning of the deposition and greatly improving reproducibility. To further optimise the spraying capabilities, we verified that most of the sprayed material was distributed in a focused area (>90% in a 1 cm radius area), thus minimising sample loss and making it particularly well-suited to the handling of valuable biological material. We set up the spray geometry to obtain a small area of spraying, corresponding to the size of the analytical surface in question, such as a 2 mm^2^ prism in the case of Fourier transform infrared spectroscopy in attenuated total reflection (FTIR-ATR). This procedure is demonstrated by the study of the distribution of droplets generated by the spray devices when a solution containing a dye (fluorescein) was sprayed onto a glass coverslip and imaged using fluorescence microscopy. The sprayed droplets exhibited a Gaussian distribution of the fluorescence intensity, with the majority of sample being deposited in a central 5 mm^2^ area and an approximate total spray radius of 1.5 cm (**Fig. S3**).

Finally, in all the following experiments, the spraying was optimised to generate a single layer of small droplets on the surface, with diameter in the 3-20 μm range which undergo fast (ms) drying time, and so that coalescence of droplets does not occur (**Supplementary Note 3**) ^1^.

### Lab-on-a-chip deposition for micro- and nanoscopic analysis over multiple length scales

Microscopic and nanoscopic methods such as EM are powerful high-throughput tools employed in most chemistry and biology laboratories for investigating biomolecular material and processes^14^. Moreover, with the recent advent of cryo-electron microscopy (cryo-EM), EM has become a standard method for the assessment and routine screenings of bio-organic samples before more detailed and higher-resolution experiments^14,15^. However, sample preparation for EM remains one of the most challenging aspects of the technique, particularly when comparison is required with samples analysed in a vitrified state by cryo-EM. Careful deposition is required to prepare samples, leaving room for possible discrepancies between individual preparations^16^. This is particularly relevant in the case of a negative stain transmission electron microscopy (TEM) experiments, where poor staining of non-conductive biological material and large stain deposits hinder high-throughput visualisation and analysis. As rinsing is required to remove excess sample and stain, the data may not reflect the full sample composition due to differential adsorption effects of heterogeneous biological material.

To overcome these types of limitations of conventional samples preparation for EM, we first demonstrate the application of our lab-on-a-chip microfluidic spray to characterise organic, biological samples and living organisms over a wide range of biological scales (**Fig. 2**). We exploit the lab-on-a-chip capabilities of our double-inlet spray device to directly stain samples on-chip (**Fig. 2a**). One channel containing sample and one channel containing uranyl acetate stain were intersected and mixed on-chip before being sprayed directly onto TEM grids. Thus, we create a single step sample preparation protocol for EM with a high degree of reproducibility between operators (**Fig. 2b**). No rinsing step was performed before imaging and analysis.

**Figure 2.**
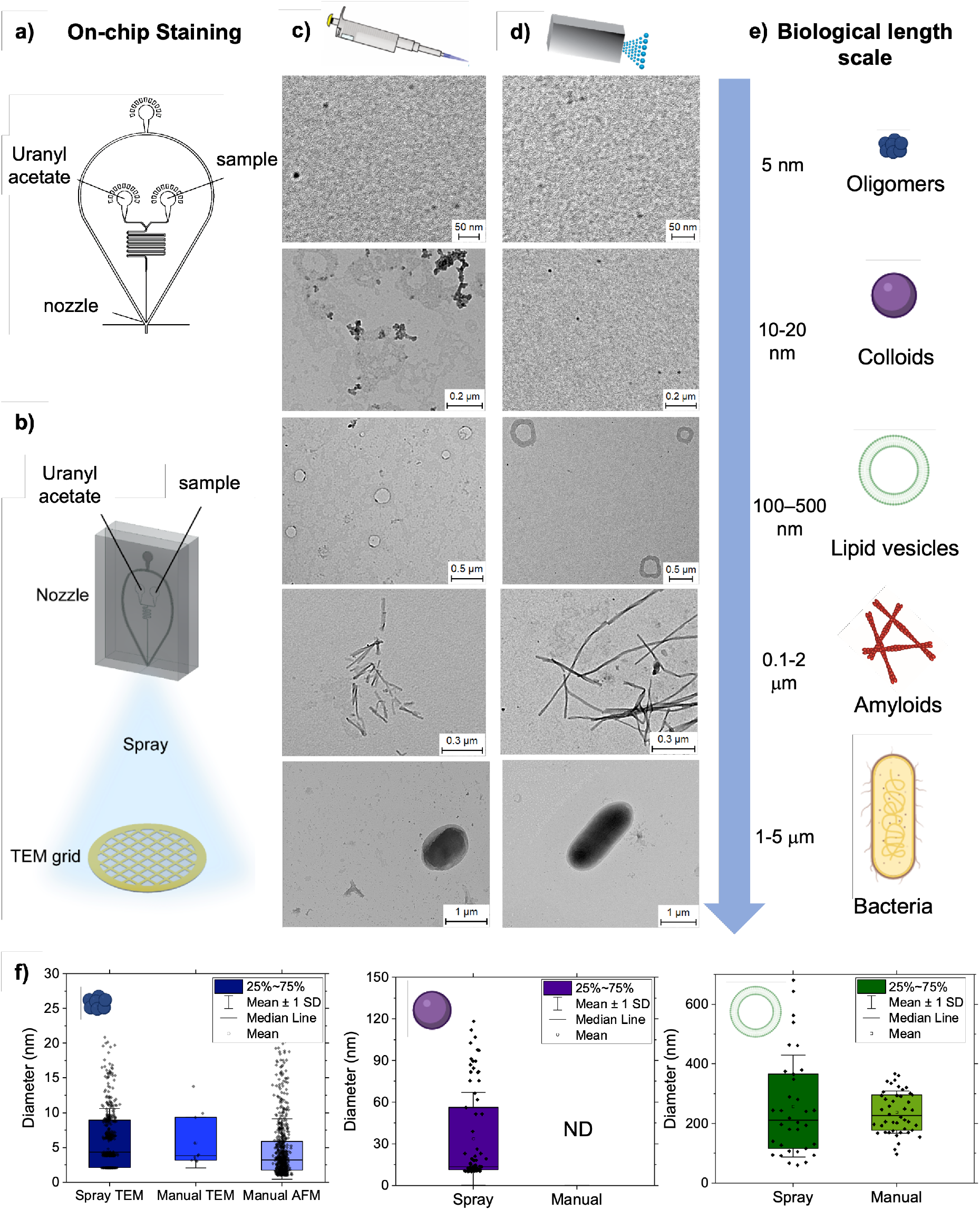
Lab-on-a-chip microfluidic deposition of biological samples over multiple biological length scales for electron microscopy. **(a, b)** A double-inlet spray nozzle was developed to allow for on-chip staining. One solution channel contained the sample, and the other contained the stain, allowing for a single-step deposition. **(c-e)** Comparison of conventional manual deposition **(c)** and lab-on-a-chip microfluidic spray deposition **(d)** for organic molecules, biomolecules and living organisms over multiple biological scales **(e). (f)** The diameter of oligomers (left), colloids (centre), and lipid vesicles (right) were measured and compared, with a larger range of sizes typically reported for samples prepared via microfluidic spray deposition, indicating that some species present in solution are not represented when deposition involves a rinsing step. Error bars represent the standard deviation (SD) of the average measured diameter for each species.

To prove the capabilities of our approach, we compared TEM measurements of samples from a few nanometers to several micrometre-scale prepared by manual deposition (**Fig. 2c**) and lab-on-a-chip microfluidic spray (**Fig. 2d**). Manual deposition was carefully performed sample deposition and staining steps (**Materials & Methods**). We considered the following species, in order of length scale (**Fig. 2e**): amyloidogenic protein oligomers (Amyloid β (Aβ), ∼5 nm), colloids formed from the aggregation of a small molecule (bexarotene, ∼10-20 nm) ^17^, lipid vesicles (phosphatidylcholine and phosphatidylserine, ∼100-500 nm), amyloid Aβ fibrils (∼0.1-2 μm), and individual bacteria (*E. coli* ∼1-5 μm).

In our hands, the manual preparation was not able to successfully prepare all samples. The colloids aggregated during drying, as previously reported ^18^. The manual deposition was also not sufficiently gentle to maintain the integrity of lipid vesicles, which often show deformation effects due to the drying process (**Fig. 2c** and **Fig. S5**) ^19,20^. The lab-on-a-chip spray deposition allowed us to successfully prepare all samples. The single-step spray deposition resulted in samples with minimal excess of uranyl acetate stain deposits on the surface, so a rinsing step was not required (**Fig. 2d**). This eliminates the effect of differential adsorption on the analysis of heterogeneous biomolecules caused by manual preparations. Thus, our lab-on-a-chip, single-step microfluidic deposition enables us to accurately study the full breadth of populations present in each heterogeneous sample.

Comparing to manual deposition methods, we see a broader size distribution of oligomers and lipid vesicles, indicating that particular species are preserved in spray deposition which are lost in manual deposition methods (**Fig. 2f**). Thus, we aimed at quantitatively demonstrating that the sample preparation is improved when deposited via lab-on-a-chip spray, compared to manual deposition. We first performed a single-molecule statistical analysis of the molecules on the surface prepared via the two methods, and then we compared these results to the measured dimensions of the molecules in solution as measured by bulk dynamic light scattering (DLS). We considered the case of the oligomeric, colloidal and lipid samples (**Fig. 2f** and **Fig. S4**). For all samples deposited via microfluidic spray, the measured size distribution corresponded well to the distributions measured in solution, while manual deposition measurements displayed a narrower size distribution, and represented less sample heterogeneity, compared to solution measurements (**Supplementary Note 4**).

Overall, we demonstrated the broad applicability of our lab-on-a-chip microfluidic spray to study biomolecules across a wide range of biological scales, up to single bacteria size, from a few nanometers to a few micrometers, preserving their morphology, integrity and heterogeneity. The spray deposition facilitated the single-molecule analysis for a few reasons. It resulted in excellent spatial separation between molecules, such as colloids which typically cluster in bulk solution due to hydrophobic effects, and minimal stain deposits; in combination, these features facilitate high-throughput imaging and analysis^21^. Finally, as we demonstrate quantitatively in the next section (**Fig. 3**) through molecular structural analysis, our approach enables us not only to preserve morphology and heterogeneity, but also the molecular conformation of the samples of interest in physiological-like conditions.

**Figure 3.**
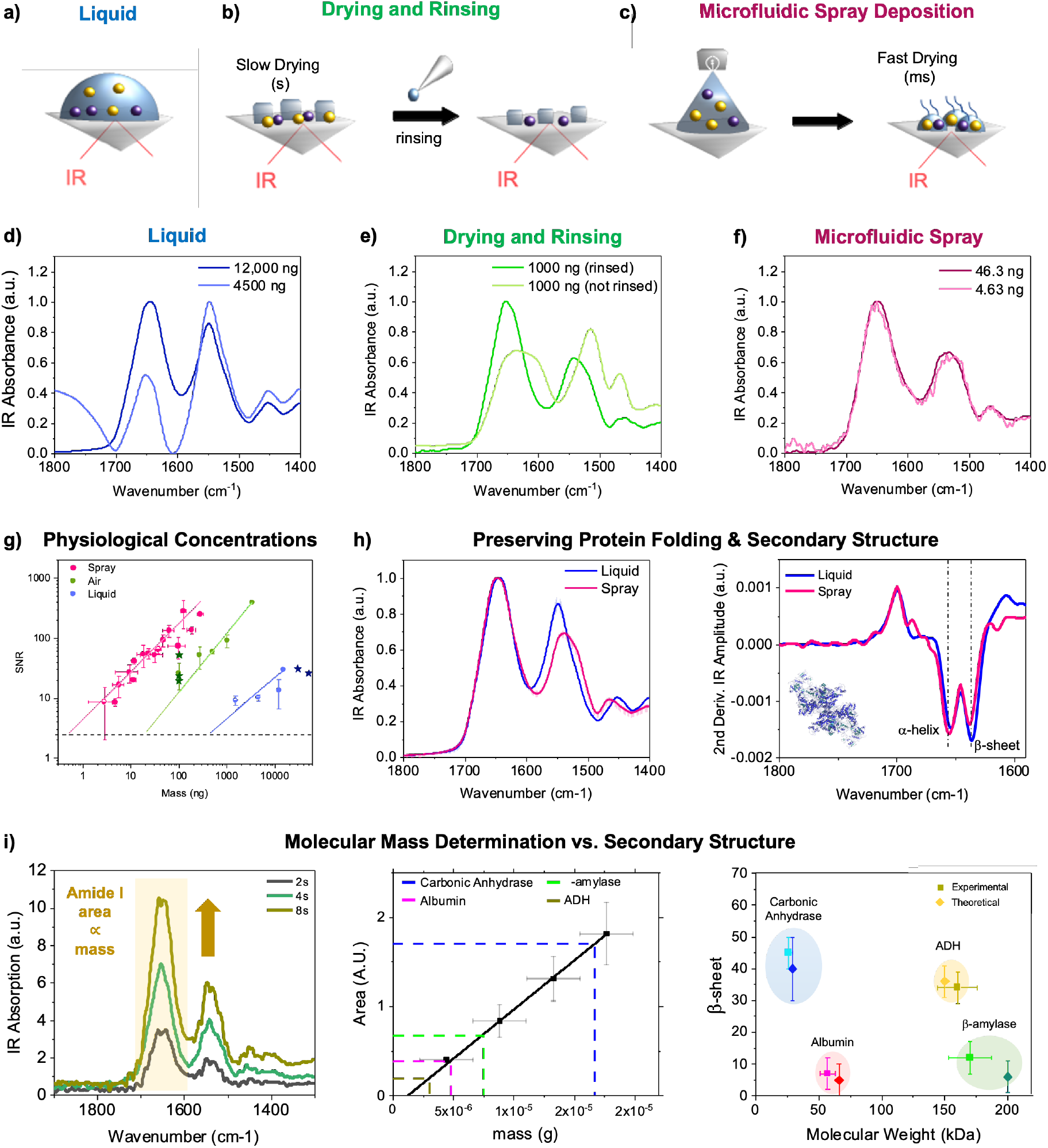
Protein identification by molecular weight and structure at physiological concentrations with nanogram sensitivity. Spectra of protein acquired by: **(a)** manual deposition of a macroscopic liquid droplet, **(b)** manual drying and rinsing of a macroscopic liquid droplet. **(c)** lab-on-a-chip microfluidic spray deposition. Spectra of protein with variable molecular weight acquired at various concentrations in liquid **(d)**, air **(e)**, and spray **(f)** (**Figs. S6-S8**). **(g)** Sensitivity in mass (ng) vs. signal-to-noise ratio (SNR) of the spectra for the different methods in **(a-c)**. The spray deposition enables the detection of nanograms amounts of protein. The SD error on the mass deposited was calculated considering a spray time error of ± 1 s (**Methods**). The sensitivity on our measurements (circles) is compared to the sensitivity of state-of-the art FTIR measurements in literature (darker stars)^23–25^. **(h)** Lab-on-a-chip microfluidic spray preserves molecular conformation as in liquid conditions (left), as demonstrated by second derivative secondary structural analysis of the amide I (right). **(i)** The area of the amide I band is proportional to the amount of protein deposited onto the ATR prism, as shown for a 11.9 µM thyroglobulin solution as a function of 3 different spray times (left). The fit of the integrated areas is used as calibration curve to determine unknown protein molecular weight (Middle panel and **Methods**). The determined molecular weight and protein secondary structure can be used for multiparameter protein identification, as shown in right panel of protein molecular weight vs. secondary structure for several proteins.

### Protein mass and secondary structure determination at nanogram concentrations

Next, we applied our lab-on-a-chip microfluidic spray for vibrational spectroscopy measurements on surfaces. Our approach enables us to perform structural and chemometric measurements of protein at low concentration in buffers, while mimicking physiological-like conditions and preserving molecular conformation (**Fig. 3)**.

Vibrational spectroscopy allows the characterisation of chemical and structural properties of biomolecules. FTIR-ATR provides high sensitivity levels at relatively low-concentrations of 1-10 mg/ml in air, and 10-100 mg/ml in liquid^2^. However, the higher concentrations needed in liquid can destabilise the biomolecules and often does not reflect their properties at physiological concentrations, while the drying process to measure in air has significant effects on the sample and spectral quality. In the case of protein solution containing stabilising buffer salts, salt crystallisation during drying can perturb the IR spectrum due to absorption and scattering effects (**Fig. S6**)^22^. To prevent these issues, reduction of the droplet size was advocated in previous studies to shorten drying times. Efforts have been made in this context to generate nL to pL droplets by means of spraying systems such as pneumatic nebulizer, thermospray and electrospray^23–28^. These efforts have resulted in spectra acquired at protein quantities as low as 1 μg, however this is in non-biological solvents or in pure water.

In order to demonstrate the novel capabilities of our device to acquire IR spectra in near-physiological conditions in a dry state, we characterised the structure and molecular weight of both globular (**Fig. 3)** and intrinsically disordered (**Fig. 4** and **Fig. S9**) proteins with different secondary structural contents. We compared the conformations of protein deposited via manual deposition in liquid (**Fig. 3a**), dried (**Fig. 3b**) and via lab-on-a-chip microfluidic spray (**Fig. 3c**).

**Figure 4.**
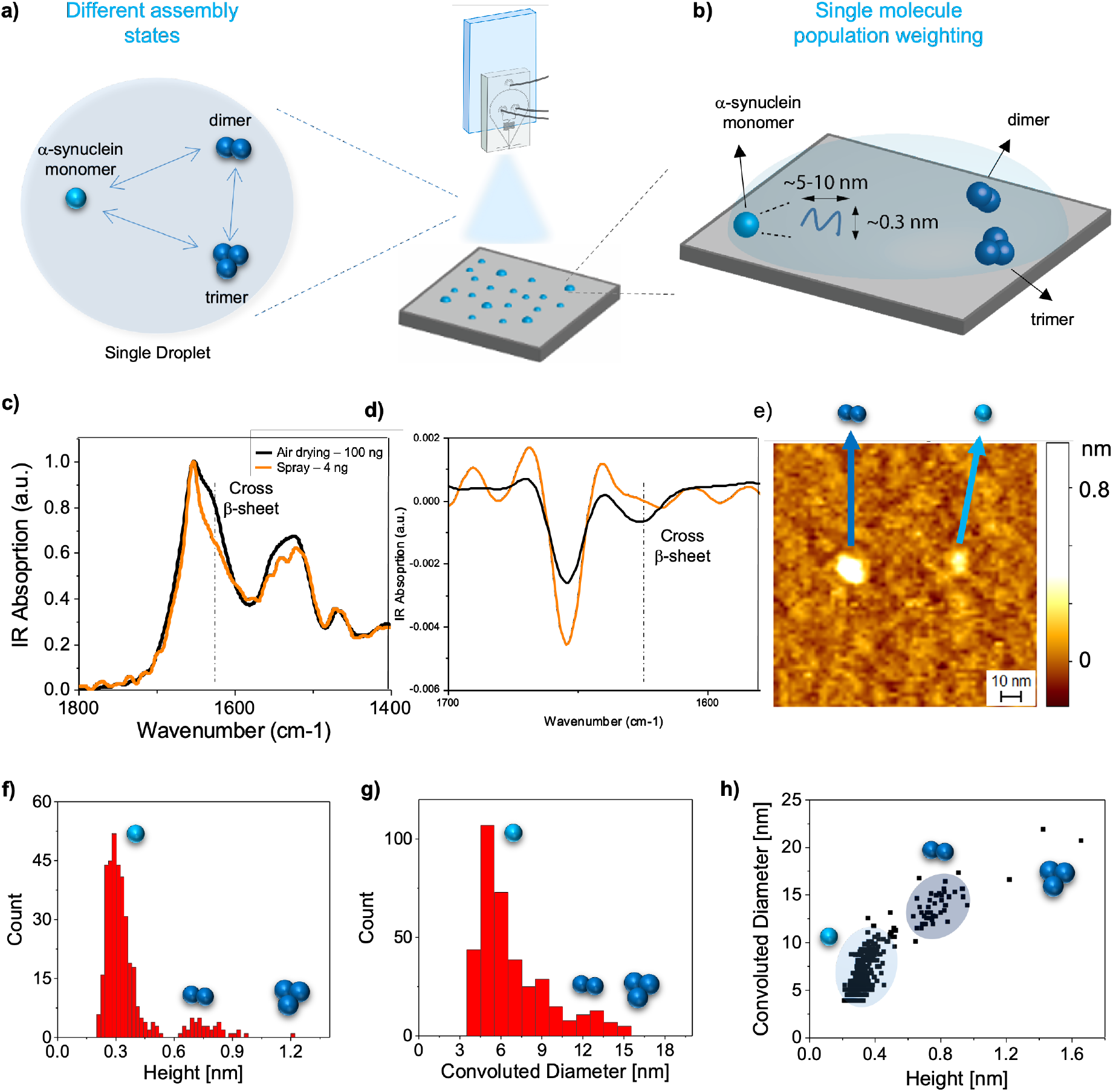
Conformational properties of the early assemblies of in an amyloid formation process. **(a)** Single-step spray deposition enables the quantitative investigation of the different self-assembled states of α-synuclein using single-molecule AFM. **(b)** Ultrafast drying better maintains the conformation of monomeric α-synuclein, compared to conventional preparation methods, as determined by ATR-FTIR. Thus, sample heterogeneity and conformations of the protein in solution can be represented on a 2D surface and imaged using AFM. **(c**,**d)** Comparison of the ATR-FTIR spectra of α-synuclein deposited via conventional air drying and via microfluidic spray deposition. A shoulder is observed at 1628 cm^-1^ for the manually deposited samples, indicating the presence of the characteristic cross-β structure of amyloid fibrils, which is not observed for the sprayed sample. This is further demonstrated by the second derivative analysis of the amide I band in (d). **(e-h)** Samples were deposited via microfluidic spray directly onto mica and imaged via AFM. Monomers, dimers and trimers can be resolved. The cross-sectional height (f) and diameter (convoluted by tip effects, g) were measured for all species. From these measurements, the populations of monomers, dimers and trimers were determined (h).

First, these measurements enabled the assessment of the sensitivity of our lab-on-a-chip approach compared to manual deposition in air and in liquid, as illustrated in the case of thyroglobulin (**Fig. 3d-f**). Spectra were reliably acquired in liquid (**Fig. 3d)** down to 12 μg of protein, but lower sample concentration resulted in significant distortions of spectra (**Supplementary Note 4**). Thus, to measure lower concentrations, FTIR spectra were acquired in air (dried samples) (**Fig. 3e**). However, the presence of buffer salts resulted in spectral distortion at high sample mass of 1 μg. Reliable spectra were acquired after a rinsing step to remove salts, which allowed us to measure spectra using 100 ng of protein. However, the rinsing step can perturb the heterogeneity and molecular conformation of the sample. Our single-step lab-on-a-chip spray deposition enabled the characterization of extremely small amounts of protein on the surface of the IR prism in the presence of salt (**Fig. 3g**), as low as 3-4 ng, while fully preserving spectral quality for structural and chemometric characterisation (**Fig. 3f-g**). We also measured the signal-noise-ratio of spectra at each concentration (**Fig. 3g** and **Methods**), demonstrating a 1000-fold increase in sensitivity when using microfluidic spray deposition compared to measurements in liquid, and 100-fold compared to traditional slow drying and rinsing. We now rival the latest advances in vibrational spectroscopies equipment and preparation methods ^23–25,29^. The improvement in sensitivity that we achieved can be rationalised by considering the reduced salt crystallisation occurring with fast drying microdroplets (**Fig. S6** and **Supplementary Note 5**).

We next sought to show that our lab-on-a-chip microfluidic spray deposition preserves the structural state of the sample during drying, similarly to when studied in a hydrated state in liquid. We compared the spectra of several globular proteins, measured in liquid at high-concentrations and when deposited by lab-on-a-chip microfluidic deposition (**Fig. 3h** left and **Figs. S6-S8**). The spray spectra presented only minor differences with the secondary structure of protein in liquid (<10%), as proved by the second derivative analysis of the shape of the amide I band (**Fig. 3h**, right). The use of globular proteins with different secondary structural components also ensures we have broad applicability to various protein folds (**Fig. S7**).

Finally, we demonstrated simultaneous molecular mass and structure determination of protein with high accuracy (**Fig. 3i** and **Methods**). By lab-on-a-chip spray, we can control the amount of protein deposited onto the surface of the prism with high levels of reproducibility by controlling the spray parameters. The area of the amide band I is directly proportional to the number of amino acids present on the surface (**Fig. 3I, left**). Thus, the area of this band will be directly proportional to the mass of protein sprayed on the prism, which can be calculated knowing the molecular weight of the protein used, flow rate, the spray time and the percentage of protein that is sprayed directly on the sensitive area of the FTIR-ATR prism (**Fig. S3**). In **Fig. 3i** central panel, we show the measured mass using a solution of thyroglobulin protein to obtain a calibration curve (black linear fit). Then, if we measure the area of the amide band I of an unknown protein, we can back-calculate its mass and molecular weight (**Methods**), thus allowing simultaneous identification of protein secondary structure and molecular weight (**Fig. 3i** right panel).

Overall, our lab-on-a-chip microfluidic spray deposition enables us to overcome the current limitations of conventional IR and vibrational methods. For the first time, we demonstrate the ability to perform simultaneous chemometric mass and secondary structure determination of unknown proteins down to the order of a single nanogram in physiological-like conditions, thus mimicking protein structure and concentrations *in vivo* ^23–29^.

### Conformational properties of the early assemblies of an amyloid formation process

Above, using the lab-on-a-chip microfluidic spray deposition with flow control, mixing capabilities and ultrafast drying, we demonstrated that it is possible to preserve both sample heterogeneity and conformation, enabling the quantitative study of protein aggregates using electron microscopy and vibrational IR bulk spectroscopy. In this section, we describe the ability of the microfluidic spray to characterise a notoriously difficult to study amyloidogenic protein sample containing heterogeneous, dynamic assembly states.

Traditional preparation methods may frustrate the quantitative study of heterogenous solutions of self-assembly of amyloidogenic proteins, because of selective adsorption effects and artificial self-assembly on surface. In this section, we demonstrate that the capabilities of our lab-on-a-chip method can be applied to quantitatively study the conformational heterogeneity of a freshly filtered and dissolved α-synuclein protein during its early molecular assembly, which is directly involved in the onset of Parkinson’s disease and neurodegeneration (**Fig. 4a,b**).

First, we used vibrational ATR-FTIR spectroscopy to acquire IR spectra and investigate the state of the aggregation-prone solution of α-synuclein when prepared by manual or lab-on-a-chip deposition (**Fig. 4c**). To assess protein secondary structure, we studied the shape of the amide I band by second derivative analysis^30^. The sample deposited by traditional drying methods showed two major peaks (**Fig. 4d**): i) the first related to α-helical conformation at 1654 cm^-1^; ii) the second related to intermolecular β-sheet content at 1625 cm^-1^; thus, demonstrating that unwanted surface-driven aggregation into amyloid species is occurring on the ATR prism. The IR spectrum of α-synuclein deposited by microfluidic spray exhibited instead only a major single peak at 1645-1654 cm^-1^ indicating the presence of random coil and α-helical conformations, which is related to the presence of monomeric and oligomeric species. Therefore, compared to manual deposition, the lab-on-a-chip microfluidic deposition maintained the conformation the solution of α-synuclein closest to its conditions in the bulk, while being able to access structural information.

Once we could demonstrate the preservation of the molecular conformation of α-synuclein in solution, we combined our lab-on-a-chip microfluidic device with the 3D high-resolution of atomic force microscopy (**Fig. 4e**, AFM), to demonstrate the ability to accurately weight heterogeneous and transient populations during the early assembly of α-synuclein. The fresh preparation of α-synuclein was deposited directly onto mica using microfluidic spray. Crucially, fast drying enables the characterization of transient assembly states, which would otherwise not be detected due to the long incubation times associated with traditional deposition.

Compared to EM, AFM enables us to measure directly the diameter and height of the early biomolecular assemblies (**Fig. 4f,g**), thus enabling us to distinguish single monomeric species from dimer, trimers and larger oligomers^31,32^. We measured the cross-sectional height and diameter of each single monomer and aggregate on the surface. We used the height to differentiate between monomers, dimers and trimers, which were around 0.3, 0.6 and 1.2 nm, respectively, as we previously demonstrated (**Fig. 4g**)^32,33^. Few oligomers with larger cross-sectional dimensions were observed. Then, we counted the different species per unit of surface and we determined the relative abundance of each species in the first minutes of the early molecular assembly. As we deposit biomolecules on the surface in a single step with no rinsing, we assume that the relative weighting of the populations on the mica surface correspond to those in solution. We could determine a ratio between monomers, dimers and trimers of about 90:8:2%. This capability is key, because the early oligomeric species are currently considered the putative species for the disease onset^34^.

## Discussion

We have presented a general lab-on-a-chip spray for the analysis of biomolecular systems on surface by a wide array of bulk and nano analytical methods (**Fig.1**). The lab-on-a-chip mixing capabilities allow us to stain and dilute samples for a wide range of single-molecule techniques (**Fig. 2**), while the standardized setup result in fine control over sample deposition, with a high level of reproducibility. Next, we applied this setup to achieve high sensitivity for chemical identification and structural characterization of protein in physiological-like conditions, with equivalent levels of sensitivity as measurements acquired with state-of-the-art methodology in non-physiological conditions in pure water (**Fig. 3**)^23–25,27,28,35^.

Overall, we have demonstrated that this approach preserves the heterogeneity and molecular conformation of transient biomolecules over multiple biological scales and in salt-containing solution, thereby more faithfully representing the physiological state of biomolecules. The ability to prepare samples for surface-based techniques in ways that maintain conformations and assemblies present in solution, while minimising the influence of salt, opens up a fruitful avenue to study biological systems which were previously inaccessible to characterisation on surfaces. We have also demonstrated the value of these capabilities to study the morphological, structural and thermodynamical equilibrium and population weighting of the transient and nanoscale sized oligomeric populations formed by α-synuclein during its self-assembly (**Fig. 4**). The study of these populations is of fundamental importance to elucidate the molecular basis of the onset of neurodegeneration.

Finally, the advances in bulk and nano-analytical chemistry capabilities presented here create the basis for future experiments, extending the experimental possibilities of surface-based methods to include notoriously difficult to study systems. One such example is that of proteins that undergo liquid-liquid phase separation, whose high level of sensitivity to their environment, such as ionic strength, has thus far prevented their characterisation using surface-based techniques. Microfluidic spray deposition enables the characterisation of ng quantities of protein in high salt conditions, thus allowing us to chemically-characterise phase separated proteins at physiologically relevant concentrations using ATR-IR, for example.

We also anticipate the possible applications to other analytical techniques, such as Raman spectroscopy or cryo-EM. In the case of cryo-EM, the fast droplet drying minimizes ice crystallisation and allows for quantitative experiments with a high level of statistical power. The extensive characterisation of the behaviour of both solvent and biomolecules when sprayed^1,12^ enables fine control over relevant processes such as droplet evaporation and film thickness, which currently represent a significant challenge^21,36,37^. Furthermore, the lab-on-a-chip capabilities, which shorten experimental timescales from minutes to seconds, combined with ultra-fast drying, can enable time-course studies of fast-reacting systems, which represents an attractive experimental approach. While we have only presented lab-on-a-chip capabilities in the context of a double-inlet, mixing device, we also note the possibility for facile methodological extension to further lab-on-a-chip capabilities, such as on-chip reactions and desalting. In addition, the ultra-fast drying essentially acts as a ‘freeze-frame’, opening the door to characterise transient phenomena in a time-dependent manner. These experimental capabilities offer the opportunity to efficiently characterise biomolecules while minimising sample use, which is particularly beneficial for precious samples that require complex preparation or are obtained from human biopsies.

## Materials and Methods

### Production of the Aβ40 and Aβ42 peptides

Expression and purification of the recombinant Aβ (M1-40) peptide (MDAEFRHDSGYEVHHQKLVFFAEDVGSNKGAIIGLMVGGVV), denoted Aβ40, and Aβ (M1-42) peptide (MDAEFRHDSGYEVHHQKLVFFAEDVGSNKGAIIGLMVGGVVIA), denoted Aβ42, were carried out as previously described^17^. Aβ40/Aβ42 was expressed in the *Escherichia coli* BL21Gold (DE3) strain (Stratagene, La Jolla, CA) and purified by sonication and dissolving the inclusion bodies in 8 M urea, followed by ion exchange in batch mode on diethylaminoethyl cellulose resin. Fractions containing Aβ40/Aβ42 were lyophilised and further purified using a Superdex 75 HR 26/60 column (GE Healthcare, Chicago, IL), and eluates were analyzed using SDS−polyacrylamide gel electrophoresis for the presence of the desired protein product. The fractions containing the recombinant peptides were combined, frozen using liquid nitrogen, and lyophilised in either 20 mM sodium phosphate buffer, pH 8, 0.2 mM EDTA (for Aβ42) or in 50 mM ammonium acetate, pH 8.5 (for Aβ40). The lyophilised product were then stored at −80 °C.

### Production of α-synuclein

Recombinant α-synuclein was expressed into BL21-competent cells and purified as described previously with slight modifications (Hoyer, JMB, 2002). Cell pellets were resuspended in 10 mM Tris, pH 8.0, 1 mM EDTA, 1 mM PMSF and lysed by multiple freeze–thaw cycles and sonication. The cell suspension was boiled for 20 min and centrifuged at 13500 rpm with a JA-20 rotor (Beckman Coulter). Streptomycin sulfate was added to the supernatant to a final concentration of 10 mg/mL and the mixture was stirred for 15 min at 4 °C. After centrifugation at 13,500 rpm, the supernatant was taken with an addition of 0.36 g/mL ammonium sulfate. The solution was stirred for 30 min at 4 °C and centrifuged again at 13,500 rpm. The pellet was resuspended and dialysed overnight against buffer containing 25 mM Tris, pH 7.7, 1 mM EDTA. Ion-exchange chromatography was then performed using a Q Sepharose HP HiScale 26/20 of buffer A (25 mM Tris, pH 7.7, 1mM EDTA) and buffer B (25 mM Tris, pH 7.7, 1 mM EDTA, 1M NaCl). The fractions containing *α*-synuclein were pooled together, and further purified using a HiLoad 26/600 Superdex 75 pg column in 20 mM NaP pH 6.5, 1 mM EDTA. α-synuclein samples were finally pooled and stored as frozen aliquots (-20 °C).

### Preparation of Aβ42 fibrils

Solutions of the lyophilised Aβ42 monomers were first dissolved in 6 M guanidinium hydrochloride, and purified from any other oligomeric species and salts using a Superdex 75 10/300 GL column (GE Healthcare, Chicago, IL) at a flow rate of 0.5 ml/min in 20 mM sodium phosphate buffer, pH 8, 0.2 mM EDTA. The centre of the peak corresponding to the monomeric protein was obtained, and the concentration was determined by absorbance using the extinction coefficient ε280 = 1490 M^−1^ cm^−1^. The obtained monomer was diluted with buffer to 5 μM, and pipetted into multiple wells of a a 96-well, half-area, black/clear flat bottom polystyrene NBS plate (Corning 3881), at 100 μl per well. Samples were then incubated at 37 °C in a plate reader (Fluostar Optima, BMG Labtech). The fibrillation process was followed by monitoring the fluorescence over time for additional Aβ42 samples that were also supplemented with 20 μM ThT. Once the fibrillation plateau was observed via fluorescence, fibrillar samples without ThT were collected from the wells into low-binding tubes.

### Preparation of Aβ40 oligomers

Aβ oligomers were prepared as previously described with slight modifications ^38^. 0.5 mg of the lyophilised Aβ40 peptide was dissolved in 300 μL of 100% HFIP overnight at 4 °C. The solvent was then evaporated under a gentle stream of N2. DMSO was then added to yield a peptide concentration of 2.2 mM, which was then sonicated twice at room temperature using a bath sonicator. The peptide was finally diluted in 20 mM sodium phosphate buffer, pH 6.9, with 200 μM ZnCl_2_ to a final concentration of 100 μM. This solution was incubated at 20 °C for 20 h, after which the oligomers were collected by centrifuging the sample at 15,000 g for 15 min at 25 °C, and resuspending the pellet in 20 mM sodium phosphate buffer, pH 6.9, with 200 μM ZnCl2.

### Preparation of small molecule colloids

Bexarotene was obtained from Sigma Aldrich at the highest purity available. Colloid formation was performed by first dissolving the bexarotene power in 10 mM solution stocks of 100% DMSO, before a serial dilution in 20 mM NaP, pH 8.0, 0.2 mM EDTA, to 100 uM in 1% DMSO (v/v) ^39^.

### Preparation of LUVs

1-palmitoyl-2-oleoyl-sn-glycero-3-phospho-L-serine (POPS) and 1-palmitoyl-2-oleoyl-sn-glycero-3-phosphocholine (POPC) were purchased from Avanti Polar Lipids, Inc. POPC and POPS in chloroform solutions were first mixed at a 7:3 lipid equivalent ratio. The solvent was then evaporated using a gentle stream of N_2_. Subsequently, POPC:POPS lipid films were dissolved in 20 mM NaP, pH 8.0, to a stock concentration of 2 mM and stirred at 45 °C for 2 h. The solution was subjected to 5 cycles of freeze-thaw, after which the preparation of large unilamellar vesicles (LUV) was done via extrusion through a 200 nm pore diameter membrane (Avanti Polar Lipids, Inc.)

### Preparation of E. coli

Glycerol stocks of previously transformed *E. coli* cells with the above recombinant Aβ42 sequence and an ampicillin resistance gene were used to inoculate overnight cultures in sterile conical flasks containing Luria–Bertani (LB) medium with ampicillin (100 μg/mL). Solutions were harvested the next day to collect the *E. coli* samples.

### Preparation of globular proteins

For FTIR measurements, proteins were dissolved in 25 and 50 mM Tris-HCl, 50 and 100 mM KCl, pH 7.4 to a concentration of 8 mg/ml. They were then filtered through a 22 μM pore and centrifuged at 22,000 g for 3 min at 4 °C, then resuspended in fresh buffer and stored at -20 °C until use. Proteins were then diluted in the same buffer to the desired concentration immediately before the experiment. All proteins were acquired from Sigma-Aldrich.

### Fabrication of microfluidic spray devices

A two-step photolithographic process was used to fabricate the master used for casting microfluidic spray devices. In brief, a 25 μm thick structure was fabricated (3025, MicroChem) was spin-coated onto a silicon wafer. This, was then soft-baked for 15 min at 95 °C. An appropriate mask was placed onto the wafer, exposed under ultraviolet light to induce polymerization, and then post-baked at 95 °C. A second 50 μm thick layer (SU-8 3050, MicroChem) was then spin-coated onto the wafer and soft-baked for 15 min at 95 °C. A second mask was then aligned with respect to the structures formed from the first mask, and the same procedure was followed, i.e. exposure to UV light and post-baking for 15 min at 95 °C. Finally, the master was developed in propylene glycol methyl ether acetate (Sigma-Aldrich) to remove any photoresist which had not cross-linked.

A 1:10 ratio of PDMS curing agent to elastomer (SYLGARD 184, Dow Corning, Midland, MI) was used to fabricate microfluidic devices. The mixture was cured for 3 hours at 65 °C. The hardened PDMS was cut and peeled off the master. The two complementary PDMS chips are then activated with O_2_ plasma (Diener Electronic, Ebhausen, Germany) and put in contact with each other and aligned precisely such that the gas inlet intersects with the liquid inlet to form a 3D nozzle ^12^.

### Use of microfluidic spray devices

Prior to introduction of sample, each device was tested and washed with MilliQ water for 5 min. Sample was then loaded into either 1 mL air-tight plastic syringes (normJet) or 200 μL air-tight glass syringes (Hamilton) and driven into the spray device using a syringe pump (Harvard apparatus). Solutions containing sample, or uranyl acetate 2% (w/v) for EM experiments, were pumped into the device with a flow rate of 100 μl/h, while the nitrogen gas inlet pressure was maintained at 3 bar. Deposition was conducted for a maximum of 10 s at a distance of 3.5 cm to ensure coalescence did not occur. Samples were sprayed directly onto the relevant surfaces (i.e. mica, FTIR prism or TEM grids) with no further washing steps required before measurements.

### Electron microscopy

Samples for TEM were prepared on 400-mesh, 3-mm copper grid carbon support film (EM Resolutions Ltd.) and stained with 2% uranyl acetate (w/v). The samples were imaged on an FEI Tecnai G2 transmission electron microscope (Cambridge Advanced Imaging Centre). Images were analyzed using the SIS Megaview II Image Capture system (Olympus). Samples for SEM/STEM were prepared using the same protocol as those for TEM. The samples were imaged on a Thermo Scientific (FEI) Talos F200X G2 TEM, operated at 30kV and 5kV for STEM and SEM respectively.

### Dynamic light scattering

All dynamic light scattering (DLS) experiments were performed with a Zetasizer Nano-S (Malvern) at 25 °C. The hydrodynamic radii were deduced from the translational diffusion coefficients using the Stokes-Einstein equation. Diffusion coefficients were inferred from the analysis of the decay of the scattered intensity autocorrelation function. All calculations were performed using the software provided by the manufacturer.

### FTIR spectroscopy

Measurements were performed on a Vertex 70 FTIR spectrometer (Bruker). Each spectrum was acquired with a scanner velocity of 20 kHz over 4000 to 400 cm^-1^ as an average of 128-412 scans. Spectra acquired in liquid were performed using a *bioATR cell II* unit and a mercury cadmium telluride (MCT) detector. Spectra acquired in air were performed using a *DiamondATR* unit and a deuterated lanthanum α-alanine-doped triglycine sulfate (DLaTGS) detector. For traditional sample deposition methods, 15 μl of sample was deposited to cover the prism. For measurements taken in liquid, spectra were acquired immediately after deposition onto the sample cell. For measurements taken in air, the sample was allowed to adsorb for 5 min, then subsequently blotted and rinsed with 10 μl MilliQ water. New background spectra were acquired before each measurement. All spectra were analysed using OPUS software (Bruker) and OriginPro (Origin Labs). Spectra presented represent 3 individual spectra which were averaged and normalised. To determine the secondary structure composition of proteins, a second derivative analysis was performed. Spectra were first smoothed by applying a Savitzky-Golay filter. In order to assess the sensitivity of each deposition method, i.e. spray, liquid and air, the signal-to-noise ratio (SNR) were calculated, both for our own data, as well as for other innovative sample deposition methods reported in the literature. The SNR was determined by dividing the intensity of the amide I peak by the RMS of the instrument noise. The SNR was then plotted versus the mass of protein deposited onto the prism (discussed below). A linear fit was applied in order to extrapolate the minimum mass required to achieve a spectrum, which is in excellent agreement with experimental data.

### Sample deposition on FTIR cell holder calculation

In order to estimate the distribution of sprayed sample arriving on the cell holder, a calibration using fluorescence microscopy was first conducted. A 0.1 mg/mL solution of a fluorescent dye, fluorescein, was sprayed onto a glass slide using the same parameters as those used in all spraying experiments, i.e. a distance of 3.5 cm with a pressure of 3 bar and a flow rate of 100 µl/h. Fluorescent images were taken at regular spatial intervals across the glass slide (**Fig. S2c-e**). By calculating the fluorescence intensity acquired from each image, the distribution of the sprayed fluorescein was plotted (blue points in **Fig. S2b**) and fitted to a Gaussian (red curve in **Fig. S2b**). Assuming that a similar distribution is observed when protein sample is sprayed onto the FTIR cell holder, we estimated the amount of protein that actually lands on the FTIR prism. As the prism has dimensions of 2 mm x 2 mm, and if the spray nozzle is placed exactly above the centre of the cell holder, then the amount of protein sample sprayed onto the holder is represented by the pink area in **Fig. S2b**. Dividing this area by the total area under the distribution, we find that 8.9% of the total protein solution sprayed lands onto the FTIR prism.

### Protein molecular weight estimation from FTIR spectra

Protein samples were sprayed onto the sample holder and their FTIR spectra were obtained. The area of spraying is accurately determined as shown in **Fig. S3**. Thus, by varying spray times for instance from 2 to 10 seconds, one can vary the amount of protein that reaches the prism. The area under the amide I band of the ATR-FTIR spectra depends on the amount of amino acids absorbing the IR light and therefore on the amount of protein on ATR-FTIR prism. Since the concentration of protein used, the molecular weight and the flow rate at which the sample was sprayed is known, and by utilising the fact that 8.9% of the total spray reaches the ATR-FTIR prism (**Fig. S3**), the mass deposited onto the FTIR holder could be calculated. We used thyroglobulin as a calibration protein with a known molecular weight (669 kDa) to generate a calibration curve, which can be described using a linear fit, as was expected (**Fig. 3**). The molecular mass of other unknown proteins was then determined using this linear relationship as calibration curve. By spraying other proteins and obtaining their FTIR spectra, the area under the amide I band was measured, by knowing the mass and volume sprayed (from the flow rate and time of sample sprayed), we could back-calculate the molecular mass of the protein under question within an error of around 11%, thanks to the calibration curve to the mass deposited in **Fig. 3g**. The error on the y-axis is derived from the area under the amide I band. It was obtained by averaging over a range of 3 spectra. The error in the mass value is a combination of 3 different factors and is shown to two standard deviations. Namely, the flow rate at which the liquid was flown through the microfluidic device, the spray deposition time, and the concentration of the thyroglobulin solution used for the calibration curve.

