## Supplementary Information for "Enhanced surface nano-analytics of transient biomolecular processes"

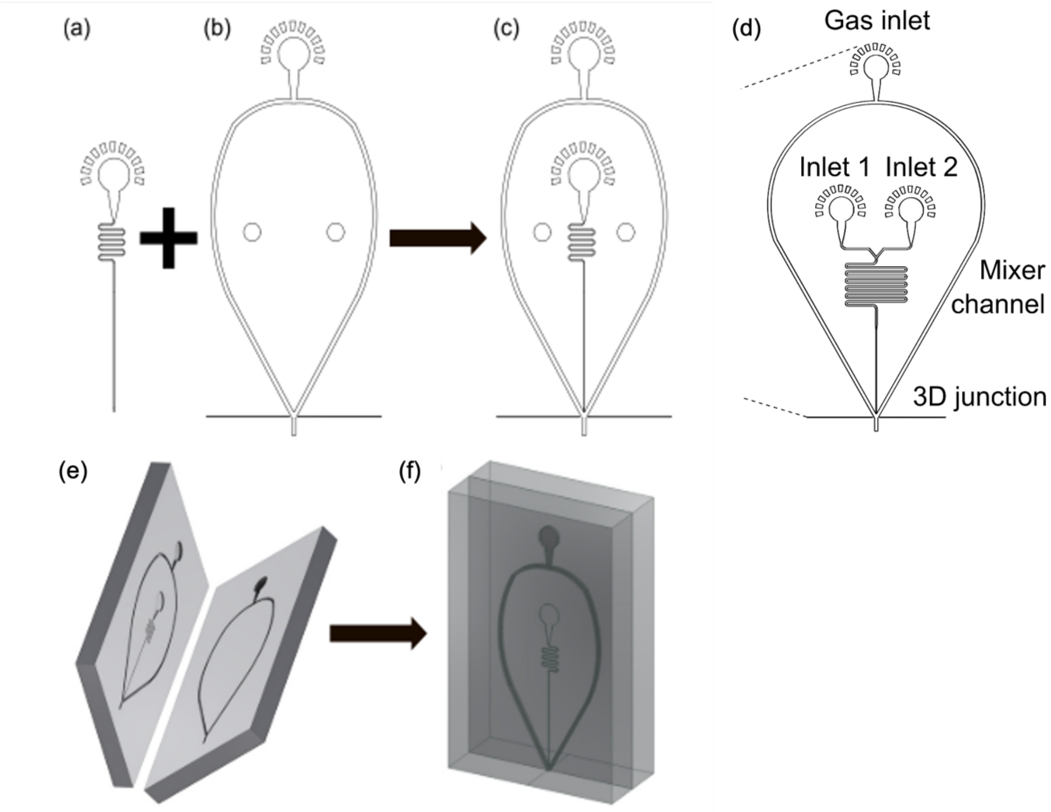


**Figure S1. Schematic of the single and double inlet microfluidic device designs**. (a-c) 25 μm high liquid channel (a), 50 μm high gas channel (b), and their combination using a two-step lithography process (c). The same applies for the double-inlet device, which simply has an additional liquid inlet and a longer channel to allow for sufficient mixing. (e-f) Assembly of the two PDMS layers occurs via plasma bonding. The PDMS pieces in (e) come from the two respective masters which are carefully aligned with respect to one another and left to bond to give the completed device (f). Adapted from ref (12).

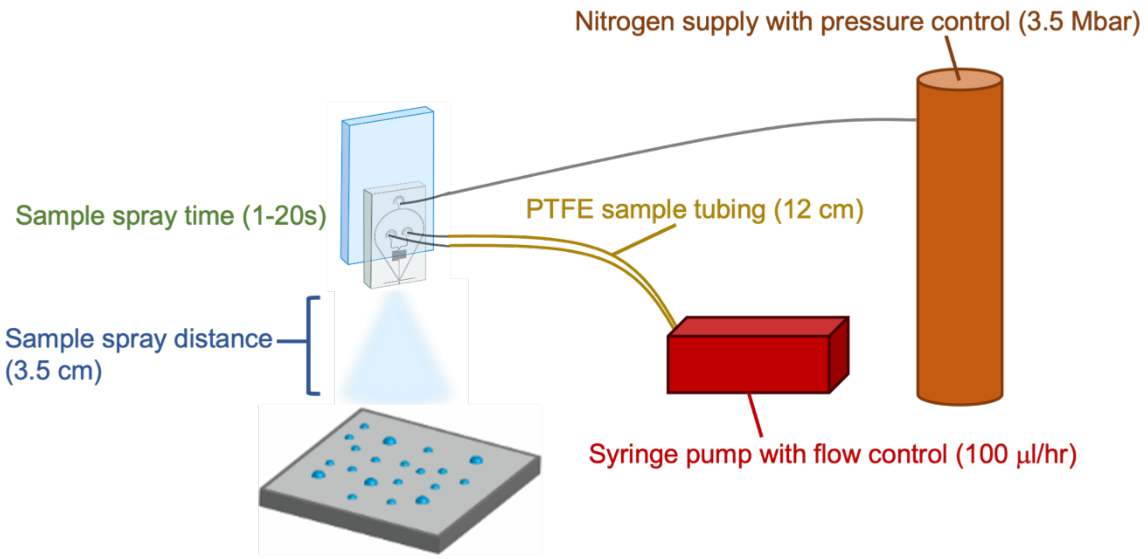


**Figure S2.** A cartoon depiction of the standardised experimental set-up. The following parameters were adjusted to enhance deposition reproducibility between experimentalists and to minimise sample usage: sample inlet flow rate, gas inlet pressure, tubing material and length, the time which sample is sprayed, and the distance between the spray device and the surface. A detailed description is included in the Methods section.

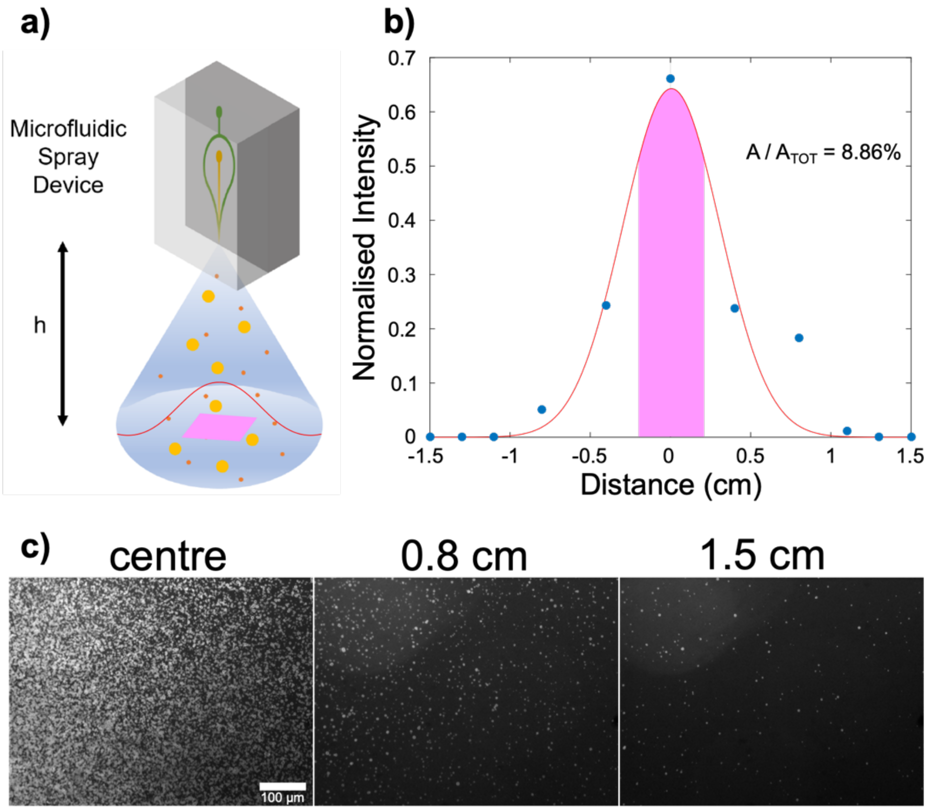

**Figure S3. Characterisation of spray generated using a microfluidic device.** (a) Schematic of the spray device. The spray area is represented in light blue, and the sample by the yellow and orange dots. (b) The distribution of the spray was generated by spraying fluorescein (1mg/ml) and calculating the fluorescence intensity at various positions (indicated by blue dots). The spray exhibits a Gaussian distribution (red line), with the majority of sample being deposited in the central 5 $mm^2$ region, as indicated by the pink area. (c) Representative fluorescence images at various distances from the spray centre show the micron-scale droplets which are generated.

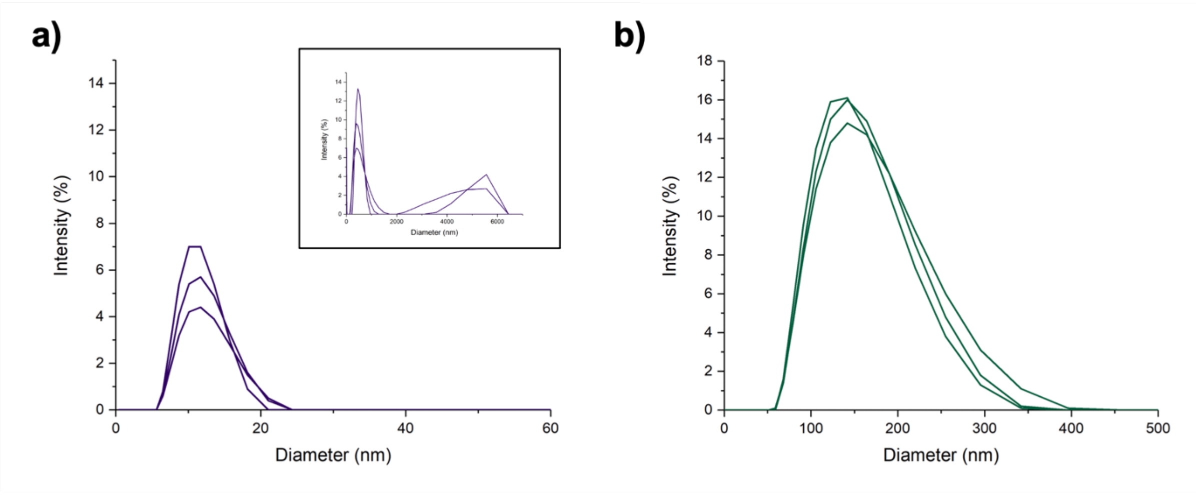


**Figure S4.** Bulk DLS data showing the diameters of colloids (a) and lipid vesicles (b) in solution. We note the presence of large aggregates present in the colloid sample.

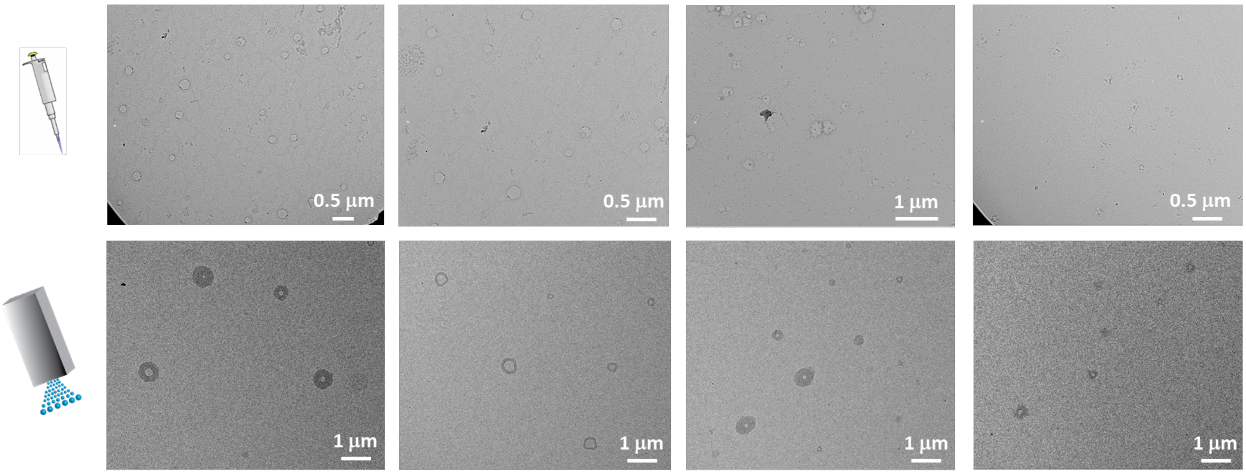


**Figure S5.** Comparison of lipid vesicles by a) manual and b) spray deposition.

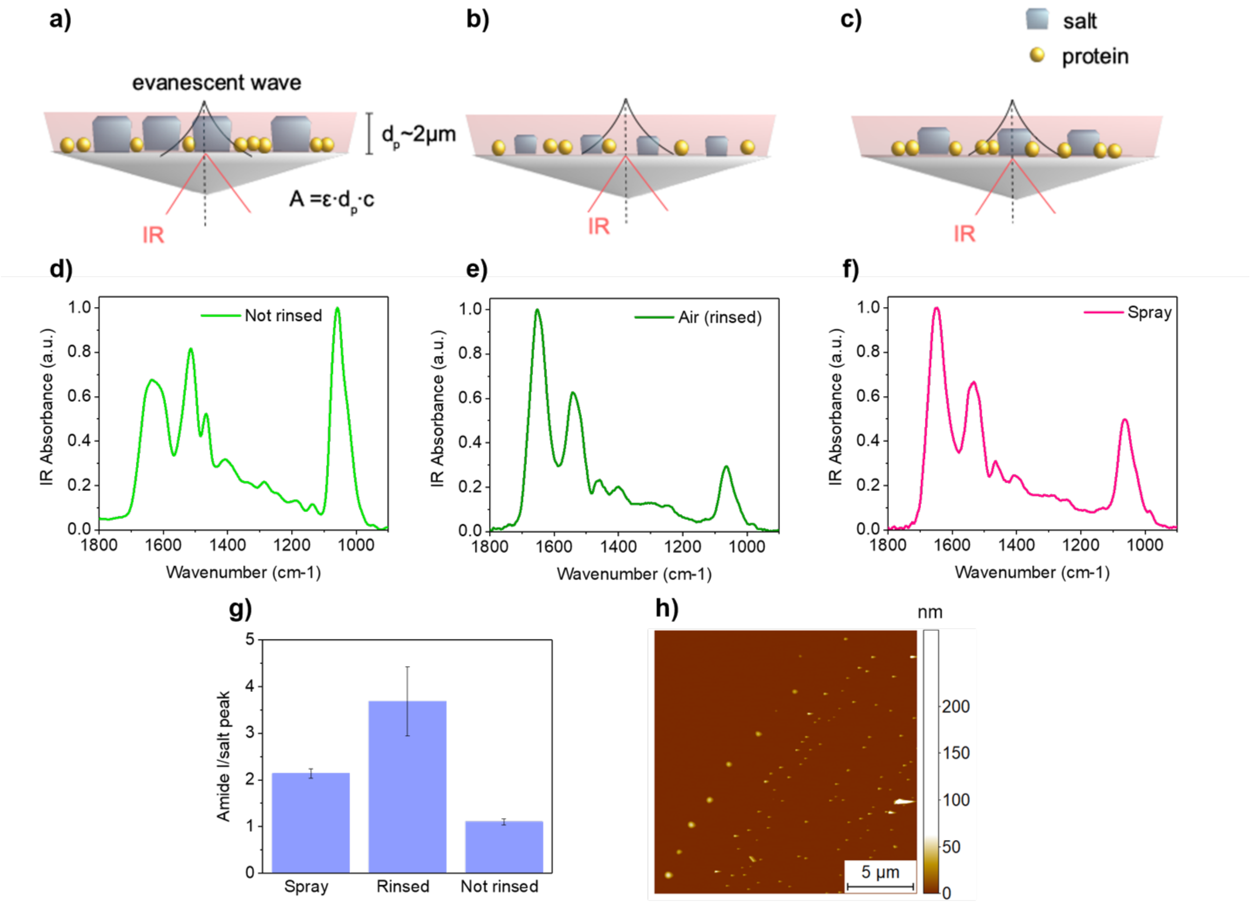


**Figure S6. Influence of salt on the IR spectra. (a-f)** Cartoon depicting the relative sizes of protein and salt on the FTIR-ATR prism in air: without rinsing (a), with rinsing (b) and sprayed (c) (not to scale), and the corresponding spectra of thyroglobulin (1 mg/ml) measured in buffer (d, e, f, respectively). (g) In order to understand the contributions of salt and protein to the IR spectra, the ratios between the area under the salt peak (~1050 $cm^{-1}$) and the amide I peak (~1650 $cm^{-1}$) were calculated for each preparation method. Error bars represent the SD. Despite containing the same ration between salt and protein, the relative influence of salt on the IR spectra is more important when deposited manually than when deposited via spray. (h) When samples dry slowly (as in traditional deposition methods), large salt crystals form which result in distorted protein spectra. As such, rinsing is required in order to remove salt crystals. In the case of microfluidic spray, the fast timescales of drying limit salt crystallisation and enable the easy measurement of proteins in buffer. Salt crystals were measured using AFM; a representative image is presented. A coffee-ring effect is observed, with salt crystals forming a ring-like shape at the edge of the droplet. The average crystal size is 95 ± 65 nm.

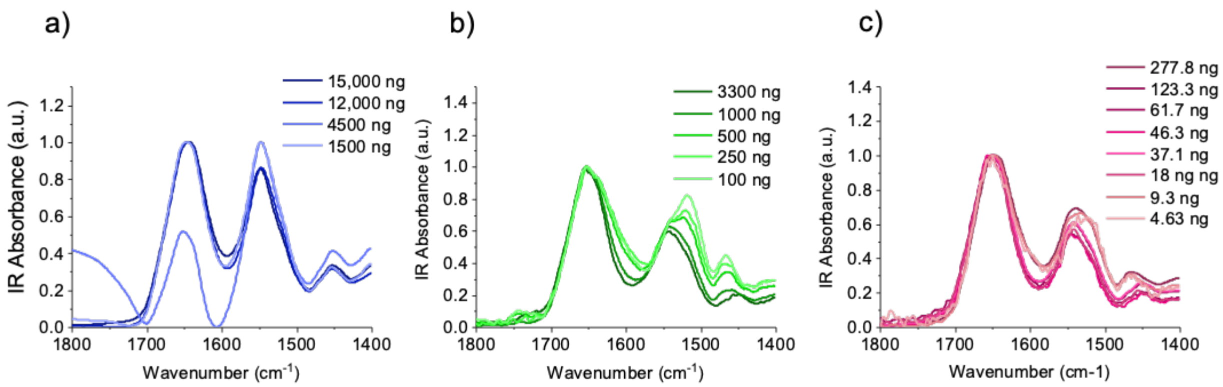


**Figure S7. Thyroglobulin IR spectra at a function of mass.** (a-c) Spectra are presented for thyroglobulin at a range of masses, for spectra acquired in liquid (a), air (b) and sprayed (c). Spectra are reproducible at a variety of concentrations when deposited via microfluidic spray. Where very small amounts of protein are deposited on the surface (<2 ng), the influence of salt becomes more apparent; secondary structure analysis (using second derivative) is still possible.

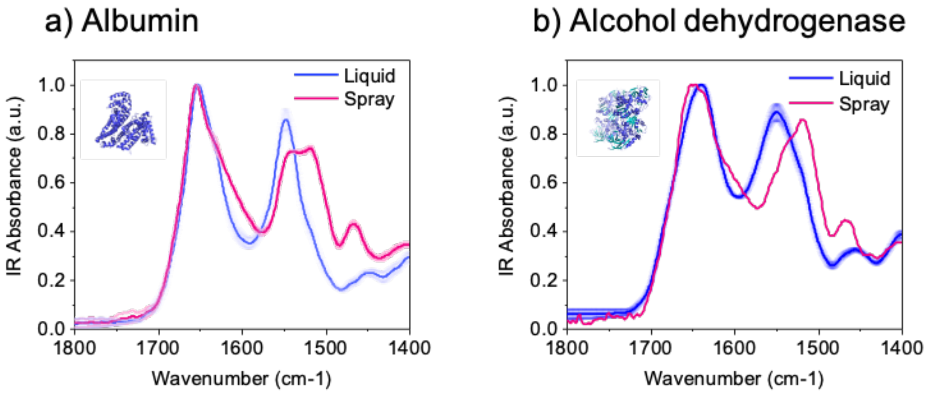

**Figure S8. FTIR spectra of albumin and alcohol dehydrogenase.** Spectra were obtained for samples deposited via microfluidic spray and in a liquid environment.

| Protein | α-helix [%] | β-sheet [%] | β-turn [%] | Coil/other [%] |
|---|---|---|---|---|
| α-synuclein | - | - | - | 100 |
| carbonic anhydrase (a) | 16.2 | 31.9 | 27.7 | 24.2 |
| β-amylase (b) | 39.6 | 12.2 | 23.1 | 25.1 |
| thyroglobulin (c) | 21.5 | 15.9 | 30.5 | 32.1 |
| albumin (d) | 70.5 | - | 9.4 | 20.1 |
| alcohol dehydrogenase (e) | 25.6 | 29.3 | 15.0 | 30.1 |

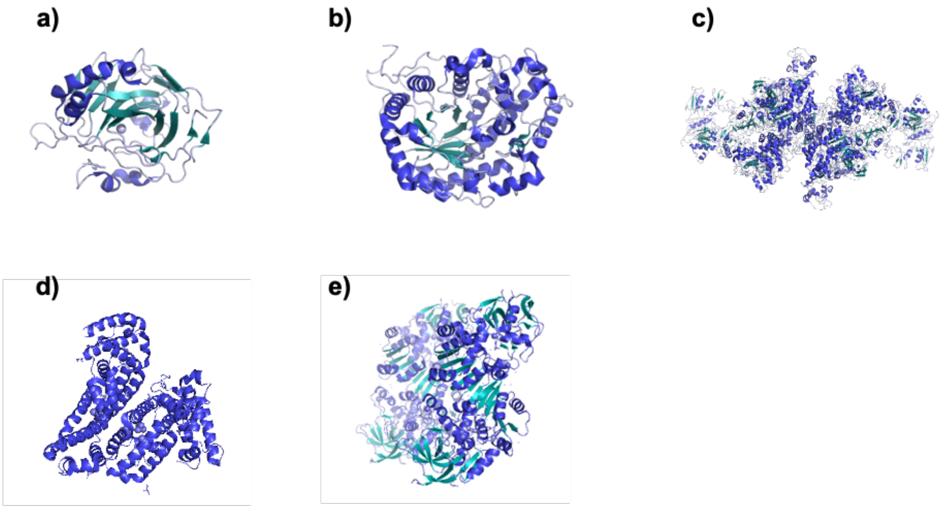


**Figure S9. Properties of globular proteins.** The secondary structure and corresponding structure of proteins are presented for the following proteins: carbonic anhydrase (5YUJ), β-amylase (1FA2), thyroglobulin (6SCJ), albumin (4F5T), alcohol dehydrogenase (46WZ).

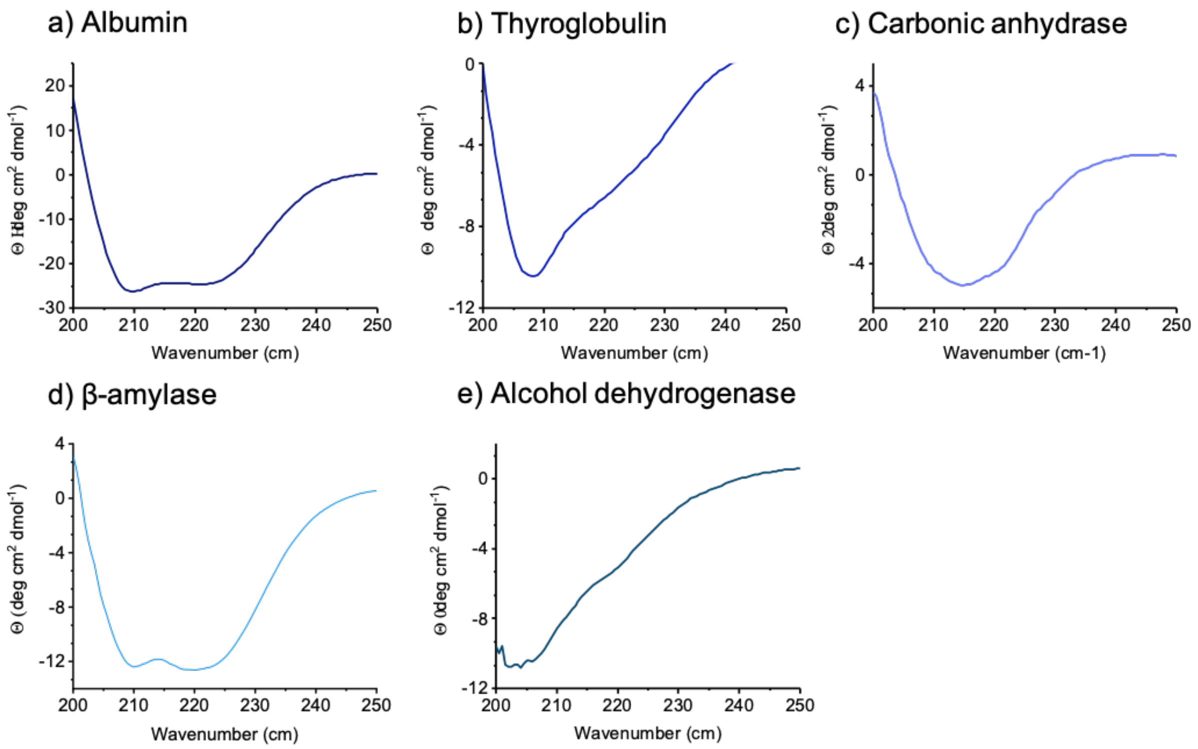


**Figure S10. CD spectra of globular proteins. (a-e)** CD spectra of globular proteins studied demonstrate that proteins are folded properly. Refs: carbonic anhydrase (40), albumin (41), β-amylase (42), alcohol dehydrogenase (43) and thyroglobulin (44).

### *SUPPLEMENTARY NOTES*

#### *Supplementary Note 1: Manual deposition of samples on surfaces*

The fixing of bio-organic and biomolecular samples on surface-based methods is often performed by manual deposition. For most analytical methods, conventional manual surface-based sample preparation involves three fundamental steps: 1) pipetting a volume of solution onto a surface and allowing it to adsorb for up to a few minutes, 2) rinsing to remove weakly adsorbed biomolecules and excess salt present in buffer, and 3) removing solvent using gentle drying. Conventionally, steps 1 and 2 are applied to measure in a liquid solution, while step 3 is necessary to measure the sample in an air or vacuum environment.

#### *Supplementary Note 2: Optimisation of microfluidic spray experimental set-up*

Every aspect of the experimental set-up was carefully considered, such as syringe size and material (**Fig. S2**). We found low volume glass syringes (250 μl) to be ideal, as this minimises sample clogging and ensures accurate volumes of liquids are flowed through the microfluidic spray device. Glass tubing proved to be incompatible with the experimental set-up due to its lack of flexibility, therefore thin polytetrafluoroethylene (PTFE) tubing was used. A standard inlet tube length of 12 cm minimised dead volumes. This allowed us to achieve a working volume for the operation of the device as low as 20 μl for each experiment, and without any minimum requirement for sample concentration.

#### *Supplementary Note 3: Droplet evaporation & salt crystallisation*

In order to understand the droplets sizes evaporation and salt crystallisation in microfluidic spray deposition, AFM images of salt crystals were analysed. The height was measured for numerous salt crystals formed, with an average height being 95 ± 65 nm. The average droplet size was measured using optical images taken using the AFM-equipped camera. Typical droplets formed were ~8-13 μm in diameter. From the droplet size, the evaporation time can be calculated (**Table S1**). This is discussed more thoroughly in ref (1).

| | **Droplet diameter (μm)** | **Evaporation time (ms)** |
|---|---|---|
| Average | 12.2 | 12.3 |
| Q1 | 8.2 | 5.5 |
| Median | 9.7 | 7.9 |
| Q2 | 12.9 | 13.8 |

**Table S1. Determination of the evaporation time of the droplets generated by microfluidic spray.** The average droplet size was measured from optical images (n=132). Note that this value is likely an overestimate, as droplets were measured using the optical camera based on the faint coffee-ring formed of salt crystals. Smaller droplets (those a few μm in diameter) will have smaller salt crystals forming that are below the detection limit of the optical camera.

According to Burton-Cabrera-Frank theory of salt crystallisation (45), from the crystal size and evaporation time, we can calculate the growth rate of salt crystals using the equation

$$G = \frac{dL}{dt}.$$

Note that height was used as a measure in crystal size instead of length ($L$). This is due to artefacts in AFM images preventing reliable measurement of the lateral dimensions of crystals. From this, we can say the growth rate of salt crystals is ~$1 \times 10^{-5}$ μm/s, which is in agreement with literature values (46–48).

Next, we applied the calculated crystal growth rate to traditional deposition methods with slow drying times. Assuming a very conservative estimate of a few minutes (180 s) drying time of a large, 10 μl droplet, the resultant crystal would be at least a few μm (1.8 μm) in size.

***Supplementary Note 3: Comparison of size distributions measured from EM***

The size distribution of samples deposited via microfluidic spray and manual deposition was compared to assess the preservation of sample heterogeneity of both methods. Comparisons were made to bulk DLS measurements acquired in solution. The size of the Aβ oligomers measured via spray was measured in the 2-20 nm range, compared to manual which displayed a 2-10 nm range. As bulk DLS measurements are often not compatible with measuring oligomers due to the heterogeneous, large amorphous assemblies which can form, we instead opted to compare these EM measurements with single-molecule AFM studies. AFM is particularly well-suited for the accurate size characterisation of oligomers due to the high resolution of the technique and the lack of staining step. AFM analysis showed a 1-25 nm diameter range, which is in excellent agreement with spray EM results (**Fig. 2f**).

Next, we sought to assess the size distribution of colloids. An analysis was not possible for manual preparation, as the colloid aggregation upon manual deposition prevented their accurate measurement. It is expected that colloids will cluster in solution to form a few large particles, which will then be over-represented in DLS measurements; this was not an issue with our spray measurements. Despite the presence of large (>1000 nm) particles observed via DLS, an excellent correspondence was observed between spray deposition EM and bulk DLS measurements, with most colloids being between 5-25 nm in diameter (Q1-Q3 11-56 nm) (**Fig. S4**).

Lastly, we assessed the size distribution of lipid vesicles, which displayed a broader range for spray deposition compared to manual, with sizes observed between 40-600 nm (Q1-Q3 116-365 nm) and 100-400 nm (Q1-Q3 177 – 296 nm), respectively (**Fig. 2f** and **Fig. S4**).

***Supplementary Note 4: Sensitivity of FTIR measurements acquired in liquid & air***

Measurements of thyroglobulin acquired in liquid (**Fig. 3d**) resulted in reliable spectra which reflect a conformationally-stable protein down to 12 μg. However, low sample concentration resulted in significant distortions of spectra due to difficulty in compensating for the IR absorption of water. Water predominantly absorbs IR light at ~3500 $cm^{-1}$ (symmetric and asymmetric O-H stretching) and at ~1600 $cm^{-1}$ (O-H bending). In particular, the IR absorption of water overlaps with the amide I band of protein, which is key for structural analysis of protein.

Thus, to measure lower concentrations, FTIR-ATR spectra are typically acquired using dried samples (**Fig. 3e**). However, the drying of salt in the buffer results in significant spectral distortion, which hampers chemometrics and structural analysis. Reliable spectra can be achieved using smaller amounts of protein by the addition of an extra rinsing step to remove excess salt, which allowed us to retrieve spectra with high quality amide band I and II peaks and with a good signal to noise ratio. However, the rinsing step can perturb the heterogeneity and molecular conformation of the sample. Our single-step lab-on-a-chip spray deposition enabled the characterization of extremely small amounts of protein in the presence of salt. The ability to measure spectra of small amounts of protein in salt, and associated improvement in sensitivity that we achieved, can be rationalised by considering the reduced salt crystallisation occurring with fast drying microdroplets, when compared to the formation of large salt crystals during the drying of a macroscopic liquid droplet (**Fig. S6** and **Supplementary Note 5**).

***Supplementary Note 5: Salt crystallisation effect on FTIR-ATR Spectra***

In **Fig. S5**, the effect of salt is described for thyroglobulin samples prepared using microfluidic spray, and those measured in air, with and without rinsing. The difference in sensitivity of ATR-FTIR by manual and spray deposition can be attributed to the formation of salt crystals.

The absorbance, and therefore the signal intensity, of FTIR-ATR measurements is related to the penetration depth ($d_p$) of the evanescent wave and it is proportional because of the Beer-Lambert law to the thickness of the material deposited on the prism. Thus, the absorbance is proportional to the height (thickness) of the material on the surface a thicker film or crystal salt on the surface will produce a stronger IR absorption.

As calculated in the **Supplementary Note 3**, the slow drying (seconds to minutes) of a macroscopic droplet causes the formation of large micrometre sized salt crystals, which have significant absorption and affect the IR measurement. Based on the growth rate of salt crystals of ~1 x $10^{-5}$ μm/s (**Supplementary Note 3**), we estimate that the salt crystals can grow as large as a 1.8 μm in the slow drying method. Therefore, it is expected that the IR spectrum would disproportionately reflect the presence of the salt. While the fast evaporation of the sprayed droplets minimises the time for salt crystallisation to occur, thus producing smaller nanometre sized crystals that do not significantly affect the IR measurements. In fact, the average crystal size is 95.11 ± 65.38 nm in height for samples deposited by spraying (**Fig. S5**). This effect is quantified by analysing the ratio between IR absorption peaks typical of the protein and the salt in solution (**Fig. S5**). In order to quantify the relative contribution of the salt to the IR absorbance spectra, the peak at ~1050 $cm^{-1}$ (the C-O bond in the primary alcohol groups of Tris was integrated and compared to that of the amide I peak We observed a ~2-fold increase of the ratio of the IR signal of the amide I and salt peak between microfluidic spray and air which was not rinsed, despite the concentration of protein to salt being the same in solution.

Thus, by significantly reducing the size of salt crystals, we thereby enable the acquisition of spectra in buffer with nanogram sensitivity without the need for rinsing (**Fig. S5**).
